# Deep Learning for Cross-Domain Spatial Transcriptomic Modeling of Tissue Repair

**DOI:** 10.64898/2026.05.13.724803

**Authors:** Tuan D. Pham

**Affiliations:** Barts and The London School of Medicine and Dentistry, Queen Mary University of London, Turner Street, E1 2AD, London, UK

**Keywords:** spatial biology, cross-domain deep learning, graph neural networks, tissue repair modeling, spatial recurrence analysis, regenerative medicine

## Abstract

Spatial transcriptomics enables investigation of tissue organization while preserving molecular and spatial information within intact tissues. However, existing computational methods primarily focus on clustering and batch integration and provide limited characterization of higher-order spatial organization and transferable tissue-state dynamics across heterogeneous biological systems. This study introduces a cross-domain spatial transcriptomic framework centered on recurrence-based latent tissue-state analysis, pathological fragmentation quantification, and transferable representation learning between wound repair and tumor-associated microenvironments. Human spatial transcriptomic datasets spanning cutaneous wound healing, oral squamous cell carcinoma, and head and neck squamous cell carcinoma were integrated within a graph-based latent embedding framework. Recurrence analysis was applied within latent transcriptomic space to characterize spatial organization and remodeling dynamics. A pathological fragmentation index quantified intra-tissue spatial disorganization from recurrence structure. The learned latent embeddings achieved a mean silhouette score of 0.79, demonstrating coherent separation of tissue states. Recurrence analysis revealed progressive restoration of spatial organization during wound remodeling, whereas tumor-associated tissues exhibited increased fragmentation and heterogeneous recurrence structure. Independent single-cell RNA-seq reference atlases demonstrated reproducible multicellular enrichment patterns within latent spatial niches. The proposed framework demonstrates that recurrence-inspired latent spatial analysis may provide biologically interpretable characterization of tissue organization and pathological remodeling across heterogeneous biological systems.

## 1 Introduction

Spatial transcriptomics has emerged as a transformative technology for investigating tissue organization by enabling simultaneous measurement of gene expression and spatial localization within intact biological specimens [1, 2]. Unlike conventional bulk transcriptomics or dissociated single-cell sequencing, spatial transcriptomics preserves the spatial architecture of tissue microenvironments, thereby allowing investigation of cell–cell interactions, tissue-state transitions, inflammatory niches, stromal remodeling, and pathological microenvironments at high molecular resolution [3]. Recent advances in spatial omics have substantially improved understanding of tumor ecosystems, tissue repair processes, developmental biology, fibrosis, and inflammatory disease.

Several computational frameworks have been developed to analyze spatial transcriptomic data using graph neural networks, manifold learning, and deep representation learning approaches. Methods such as SpaGCN [4], STAGATE [5], SEDR [6], DeepST [7], and GraphST [8] have demonstrated the utility of graph-based learning for identifying spatial domains, integrating transcriptomic and spatial information, and improving tissue-state clustering. More recent studies have additionally explored cross-sample integration and batch correction strategies using approaches including PRECAST [9], Harmony-based integration [10], and graph-domain adaptation methods [11, 12]. These frameworks have contributed substantially toward improving latent representation learning and spatial domain identification across heterogeneous spatial transcriptomic datasets [13].

Despite these advances, several important limitations remain. Most existing methods focus primarily on spatial clustering or batch integration within relatively homogeneous biological systems, often emphasizing tumor microenvironment analysis or single-dataset spatial organization [4, 5, 6, 8, 9]. Relatively few studies have investigated transferable cross-domain spatial representation learning across fundamentally distinct biological processes such as tissue repair and pathological remodeling [13, 14]. Furthermore, most current approaches provide limited characterization of dynamic spatial organization, tissue-state transitions, or latent structural heterogeneity within spatial transcriptomic embeddings [5, 8, 9].

Biological tissue repair is inherently spatial, heterogeneous, and temporally dynamic. Processes such as inflammation, hypoxia, extracellular matrix remodeling, fibrovascular activation, epithelial regeneration, fibrosis, and osteogenic repair evolve continuously across spatial and temporal scales. Similar biological programs are also partially recapitulated within pathological microenvironments including cancer, chronic inflammation, fibrosis, and impaired wound healing. The long-standing concept that tumors resemble “wounds that do not heal” [15] highlights the biological overlap between tissue repair and tumor-associated remodeling processes. However, computational frameworks capable of jointly modeling these related spatial biological systems remain limited.

Understanding spatial organization during tissue repair has important translational implications for regenerative medicine, surgery, oncology, and craniofacial reconstruction. In facial trauma, burn injury, chronic wounds, fibrosis, and tumor-associated remodeling, clinical outcomes are strongly influenced by localized biological microenvironments that cannot be fully characterized using conventional imaging or histopathological assessment alone [16]. Spatial transcriptomic modeling therefore provides a potential framework for identifying latent biological repair states associated with regenerative success, persistent inflammation, impaired vascularization, fibrosis progression, or pathological remodeling.

Recent spatial transcriptomic studies have characterized wound-healing dynamics, inflammatory signaling, fibroblast heterogeneity, immune activation, and extracellular matrix remodeling during tissue repair [17, 18, 19]. Similarly, spatial oncology studies have investigated tumor invasion fronts, immune exclusion, stromal activation, angiogenic niches, and hypoxic tumor microenvironments [20]. However, most existing studies remain descriptive and dataset-specific, with relatively limited emphasis on transferable latent biological representation learning across related tissue systems.

Another important limitation of current spatial transcriptomic analysis approaches is the relative lack of methods for quantifying higher-order spatial organization and tissue-state coherence within latent embedding spaces. Spatial recurrence analysis, recurrence networks, and fragmentation-based approaches have been widely applied in nonlinear dynamical systems, physiological signal analysis, and complex systems modeling [21, 22, 23], but remain relatively underexplored in spatial omics research. Such approaches may provide additional insight into spatial tissue organization, architectural stability, biological heterogeneity, and pathological fragmentation within tissue microenvironments.

The present study introduces a deep learning framework for cross-domain spatial transcriptomic representation learning of tissue repair and pathological microenvironments. The proposed methodology integrates graph neural networks, domain-aware latent representation learning, deep embedded clustering, recurrence-based spatial organization analysis, pathway-level biological scoring, and single-cell-guided cell-state projection within a unified computational framework. Unlike many existing spatial transcriptomic methods that focus primarily on clustering or batch integration, the proposed approach aims to characterize transferable latent tissue states shared across wound-healing and tumor-associated biological systems while simultaneously preserving biologically meaningful inter-domain variability.

A central component of the framework is the introduction of recurrence-inspired spatial organization analysis within latent embedding space. Recurrence matrices, local coherence maps, and pathological fragmentation indices were used to characterize spatial stability, tissue-state organization, and biological heterogeneity across wound-healing stages and pathological microenvironments. In addition, a spatial composite score integrating inflammation, hypoxia, extracellular matrix remodeling, and osteogenic activity was developed to provide biologically interpretable spatial characterization of adverse tissue microenvironments.

The framework was evaluated using human spatial transcriptomic datasets spanning cutaneous wound healing, oral squamous cell carcinoma (OSCC), and head and neck squamous cell carcinoma (HNSCC). Independent single-cell RNA sequencing reference atlases were further projected onto the learned spatial embeddings to biologically validate the inferred latent tissue states and multicellular spatial organization.

The proposed approach aims to provide a computational foundation for transferable spatial modeling of tissue repair and pathological remodeling across heterogeneous biological systems. More broadly, the framework may contribute toward future development of multimodal spatial foundation models capable of integrating spatial transcriptomics with histopathology, radiological imaging, physiological measurements, and clinical metadata for precision regenerative medicine, surgical planning, and spatial oncology.

## 2 Methods

### 2.1 Datasets

Publicly available human spatial transcriptomic and scRNA-seq datasets were used to model tissue repair processes within a cross-domain framework. Here, cross-domain modeling refers to learning from multiple datasets with different biological contexts, acquisition conditions, and tissue sources while projecting them into a shared latent representation. Table 1 summarizes the datasets used in this study and their respective roles within the computational framework.

**Table 1:** Summary of datasets and their roles in the study.

| Dataset | Type | Usage | Purpose |
| --- | --- | --- | --- |
| GSE241124 | Human spatial transcriptomics | Training and internal validation | Provides temporally resolved wound-healing states (intact, day 1, day 7, day 30) for supervised learning of tissue repair dynamics |
| GSE208253 | Human spatial transcriptomics | Cross-domain modeling and external validation | Provides heterogeneous pathological tissue microenvironments for evaluating transferable spatial tissue representations |
| GSE181300 | Human spatial transcriptomics | Cross-domain modeling and external validation | Provides an independent spatial transcriptomic domain for robustness assessment of learned tissue-state embeddings |
| GSE215403 | Human single-cell RNA-seq | Cell-state projection and biological validation | Provides single-cell reference profiles for projection of epithelial, fibroblast, endothelial, stromal, and immune-cell states onto spatial tissue regions |
| GSE181919 | Human single-cell RNA-seq | Cell-state projection and biological validation | Provides an independent single-cell reference atlas for validation of inferred cellular compositions across tissue states |

### 2.1.1 Spatial transcriptomic datasets

Three human spatial transcriptomic datasets were included. GSE241124 [17] was used as the primary labeled dataset because it contains human skin wound-healing samples across multiple temporal stages, including intact tissue, day 1, day 7, and day 30 post-injury. These temporal labels were used for supervised learning of wound-healing states. Although the tissue source is skin, the underlying biological phases of inflammation, proliferation, angiogenesis, extracellular matrix remodeling, and tissue maturation are broadly relevant to tissue repair processes across multiple anatomical systems.

GSE208253 [20] contains human oral squamous cell carcinoma (OSCC) spatial transcriptomic samples and was used as an oral-domain spatial dataset. GSE181300 [24] contains human head and neck squamous cell carcinoma (HNSCC) spatial transcriptomic samples and was used as an independent heterogeneous tissue-domain dataset. These two datasets were not assigned wound-stage labels and contributed to unsupervised cross-domain representation learning, clustering, recurrence analysis, and external spatial evaluation.

In total, 36 spatial samples were processed. For GSE241124, expression matrices were loaded from `filtered_feature_bc_matrix.h5` files and matched with spatial coordinate files extracted from the corresponding spatial image folders. For GSE208253 and GSE181300, expression matrices were loaded from `filtered_feature_bc_matrix.h5` files and matched with `tissue_positions_list.csv` and `scalefactors_json.json` files.

#### 2.1.2 Single-cell RNA sequencing datasets

Two human scRNA-seq datasets were prepared for post hoc cell-state projection: GSE215403 [25] and GSE181919 [26]. GSE215403 was available as `matrix.mtx, features.tsv`, and `barcodes.tsv` files. GSE181919 was available as UMI (Unique Molecular Identifier) count and barcode metadata tables.

### 2.2 Data Representation and Gene Harmonization

For spatial dataset *d*, the raw count matrix is denoted by

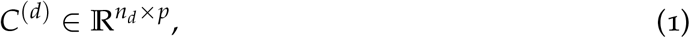

where *n*_*d*_ is the number of spatial spots and *p* is the number of genes. The corresponding spatial coordinates are

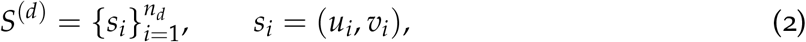

where (*u*_*i*_, *v*_*i*_) is the two-dimensional location of spot *i*.

Genes were harmonized across datasets using Ensembl gene identifiers (https://www.ensembl.org/info/genome/stable_ids/index.html), due to inconsistencies in gene symbol annotations. A mapping from Ensembl identifiers to HUGO Gene Nomenclature Committee (HGNC) gene symbols (https://www.genenames.org/) was derived from the *Homo sapiens* GRCh38 release 109 GTF (Gene Transfer Format) annotation, which provides genomic coordinates and structural information for annotated genes and transcripts. Highly variable genes were identified by ranking genes according to their variance across spatial spots, and the top 1000 genes were retained. In addition, 29 Ensembl identifiers corresponding to curated inflammatory, hypoxic, extracellular matrix, and osteogenic marker genes were forcibly retained, yielding a final feature set of 1015 genes.

The retained marker genes enabled pathway-level interpretation after mapping Ensembl identifiers back to HGNC symbols. Detected markers included *ALPL, COL1A1, CXCL8, IL1B, RUNX2, TNF*, and *VEGFA*.

### 2.3 Normalization and Standardization

Raw counts were library-size normalized and log-transformed as

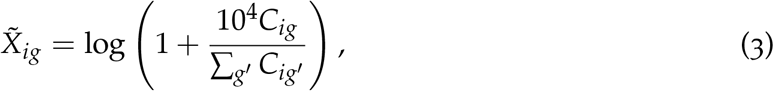

where *C*_*ig*_ is the raw count for gene *g* at spot *i*, and ∑_*g*_′ *C*_*ig*_′ is the total count at that spot. The scaling factor 10^4^ standardizes sequencing depth across spots.

The data were then standardized gene-wise as

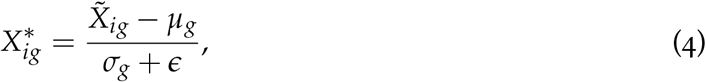

where *µ*_*g*_ and *σ*_*g*_ are the mean and standard deviation of gene *g*, and *ϵ* is a small constant used to avoid division by zero.

### 2.4 Spatial Graph Construction

Each tissue section was represented as a spatial graph as

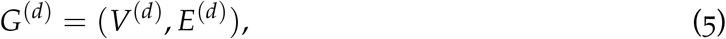

where nodes *V* correspond to spatial spots and edges *E* connect neighboring spots. Neighborhoods were defined using *k*-nearest neighbors with *k* = 6:

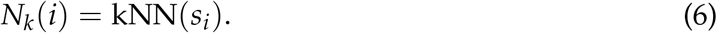

A weighted adjacency matrix was constructed using a Gaussian spatial kernel:

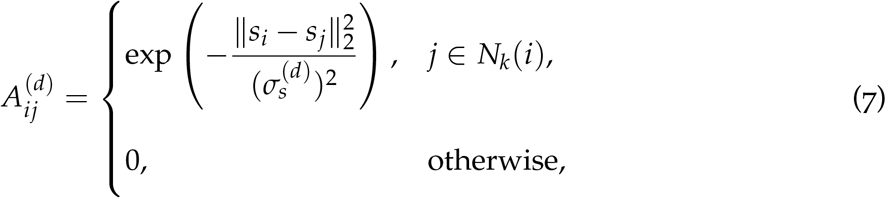

where ∥*s*_*i*_ − *s*_*j*_∥_2_ is the Euclidean distance between spots and 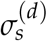 is estimated as the median neighbor distance in dataset *d*:

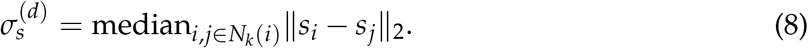

Self-loops were added and the adjacency matrix was symmetrically normalized as

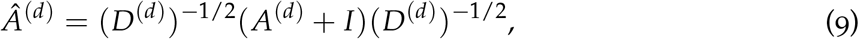

where *D*^(*d*)^ is the degree matrix and *I* is the identity matrix.

### 2.5 Graph Neural Network Encoder

The model learns a spatially informed embedding:

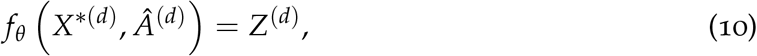

where 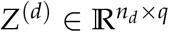 and *q* = 64 is the latent embedding dimension.

The input layer is

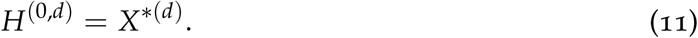

Graph convolutional propagation is defined as

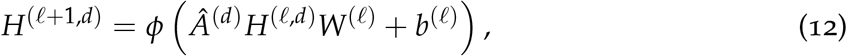

where *H*^(*ℓ,d*)^ is the layer-*ℓ* representation, *W*^(*ℓ*)^ and *b*^(*ℓ*)^ are learnable parameters, and *ϕ*(*·*) is the ReLU activation function. The encoder comprised successive layers of sizes *p*′, 256, 128, and 64, where *p*′ = 1015 denotes the number of retained genes. The final embedding is

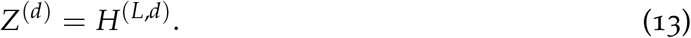

Weights were initialized using scaled Gaussian initialization:

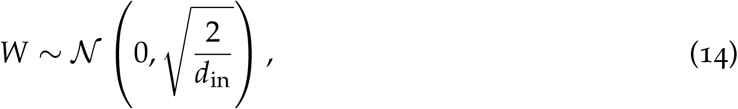

where *d*_in_ is the input dimension of the layer. Biases were initialized to zero.

### 2.6 Latent Representation Supervision Using Wound-Healing Stages

For labeled GSE241124 samples, wound-stage probabilities were predicted from the latent embedding as

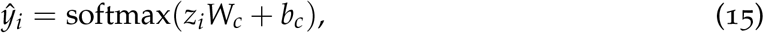

where *z*_*i*_ ∈ ℝ^64^ is the embedding of spot *i, W*_*c*_ ∈ ℝ^64*×*4^ and *b*_*c*_ ∈ ℝ^4^ are learnable classifier parameters, and the four classes correspond to intact tissue, day 1, day 7, and day 30.

The classification loss is defined as

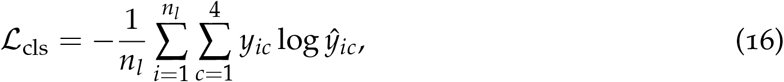

where *n*_*l*_ is the number of labeled spots and *y*_*ic*_ is the one-hot class label.

### 2.7 Domain Classification

Dataset membership was predicted using a domain classifier:

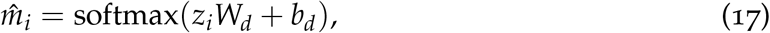

where 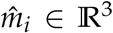 is the predicted domain probability vector, *W*_*d*_ ∈ ℝ^64*×*3^ and *b*_*d*_ ∈ ℝ^3^ are learnable parameters, and the three domains correspond to GSE241124, GSE208253, and GSE181300.

The domain classification loss is defined as

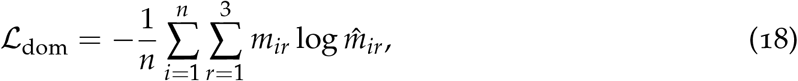

where *m*_*ir*_ is a binary indicator equal to one if spot *i* belongs to domain *r*, and zero otherwise. This loss was included to retain information about dataset structure during cross-domain representation learning.

### 2.8 Deep Embedded Clustering

Spatial tissue states were inferred using a Deep Embedded Clustering (DEC) framework [27], in which cluster assignments are learned jointly with the latent representation. This enables the embedding space to be explicitly structured into biologically meaningful groups.

Let *z*_*i*_ ∈ ℝ^*q*^ denote the latent embedding associated with spatial location *i*. Soft cluster assignments are defined using a Student’s *t*-distribution kernel with degrees of freedom fixed to *α* = 1:

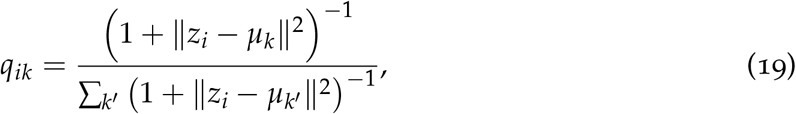

where *µ*_*k*_ ∈ ℝ^*q*^ denotes the centroid of cluster *k*. This heavy-tailed kernel assigns higher probability to nearby centroids while improving robustness to noise and promoting separation between clusters.

To refine cluster assignments, an auxiliary target distribution is constructed to emphasize high-confidence assignments:

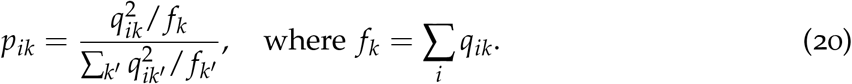

This self-training mechanism amplifies confident assignments and normalizes cluster contributions, preventing large clusters from dominating the learning process.

The clustering objective is defined as the Kullback–Leibler (KL) divergence between the target distribution *P* and the soft assignments *Q*:

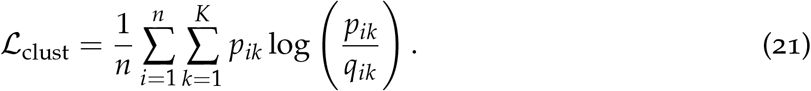

Here, KL(*P* ∥ *Q*) denotes the Kullback–Leibler divergence between the sharpened target distribution *P* = *{p*_*ik*_*}* and the current soft cluster assignment distribution *Q* = *{q*_*ik*_*}*. Minimizing KL(*P* ∥ *Q*) enables the soft assignments *Q* to match the sharpened target distribution *P*, resulting in more compact and well-separated clusters in the latent space.

The number of clusters was fixed to *K* = 5, corresponding to major tissue states associated with tissue repair, including inflammatory, fibrovascular, epithelial/interface, remodeling, and pathological niches. Each cluster is represented by a centroid *µ*_*k*_ ∈ ℝ^*q*^, with *k* = 1, …, *K*.

Training was performed in two stages. First, the encoder was pretrained for 20 warmup epochs without the clustering objective. Cluster centroids were then initialized using *k*-means on the learned embeddings. Subsequently, the clustering loss was activated, and both network parameters and cluster centroids were updated jointly via backpropagation. In contrast to the original DEC formulation [27], which operates on independent samples, the proposed framework integrates the clustering objective within a spatial graph neural network and a multi-domain learning setting. This enables the identification of spatially coherent and biologically interpretable tissue niches across heterogeneous transcriptomic datasets.

### 2.9 Joint Optimization

The total training objective is defined as

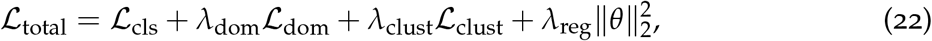

where *θ* denotes all learnable parameters. The weighting parameters were defined as: *λ*_dom_ = 0.2, *λ*_clust_ = 0.5, and *λ*_reg_ = 10^−4^. The model was trained for 100 epochs using the Adam optimizer with learning rate 10^−3^.

### 2.10 Spatial Recurrence Analysis

Following the concept of recurrence analysis in nonlinear dynamical systems [21], similarity between latent tissue states was quantified using a Gaussian recurrence matrix:

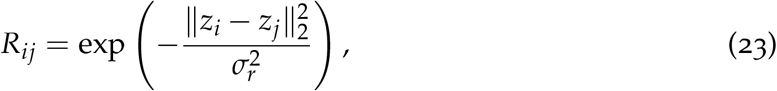

where *R*_*ij*_ is the recurrence similarity between spots *i* and *j*, and *σ*_*r*_ is estimated as the median non-zero pairwise distance between embeddings.

Local coherence quantifies the average recurrence similarity between a spatial location and its neighboring tissue regions, thereby measuring the degree of local spatial organization within the latent embedding space. Local coherence, denoted as *ρ*_*i*_, is defined as

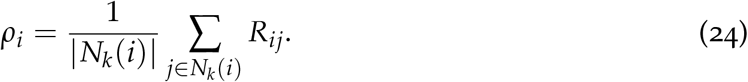

Here, *N*_*k*_(*i*) denotes the set of *k* nearest neighboring spatial locations of spot *i*, and |*N*_*k*_(*i*)| is the number of neighboring locations. Higher values of *ρ*_*i*_ indicate stronger local spatial coherence and greater similarity of neighboring tissue states within the latent embedding space, whereas low *ρ*_*i*_ indicates spatial heterogeneity.

The pathological fragmentation index (PFI) is introduced in this study as a spatial graph-based measure of intra-cluster tissue disorganization. The metric quantifies the degree to which neighboring spatial locations within a cluster exhibit heterogeneous latent biological states.

For a selected cluster *k*, the PFI is computed as

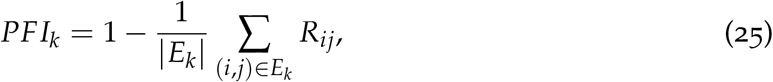

where *E*_*k*_ is the set of graph edges connecting neighboring spots assigned to cluster *k*. A high value indicates greater spatial fragmentation and reduced local consistency.

### 2.11 Biological Pathway and Spatial Composite Scoring

Pathway activity was computed using curated marker gene sets after mapping retained Ensembl IDs to HGNC symbols. For a pathway gene set *G*, the pathway score at spatial location *i* is defined as

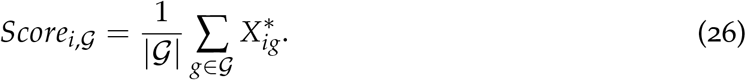

Pathway groups included inflammation, hypoxia, extracellular matrix (ECM) remodeling, angiogenesis, and osteogenesis. To summarize the functional state of each spatial location, a spatial composite score (SCS) is introduced here as a spatially resolved composite score integrating pathway activities associated with inflammation, hypoxia, extracellular matrix remodeling, and impaired osteogenic activity. The SCS is mathematically expressed as

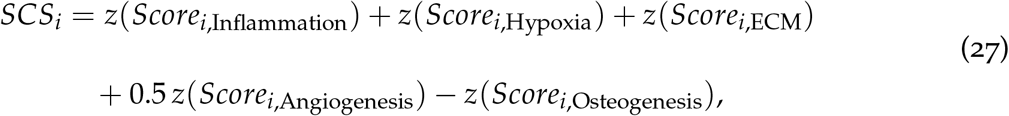

where *z*(*·*) denotes z-score standardization applied across spatial locations for each pathway. By construction, each standardized component has zero mean and unit variance across locations. Higher values of *SCS*_*i*_ indicate regions with elevated inflammatory, hypoxic, and matrix remodeling activity relative to osteogenic activity, reflecting stressed or pathological tissue states. Conversely, lower values correspond to regions with stronger osteogenic activity, indicative of reparative or regenerative processes. The angiogenesis component is assigned a reduced weight of 0.5 to reflect its context-dependent role in both physiological repair and pathological processes, thereby moderating its influence relative to more directly discriminative pathways such as inflammation and hypoxia.

### 2.12 Cell-State Projection

To infer the cellular composition at each spatial location, spatial transcriptomic profiles were projected onto a reference atlas of cell states derived from scRNA-seq. This formulation models each spatial expression profile as a convex combination of reference cell-state signatures, enabling biologically interpretable deconvolution of mixed signals.

Let 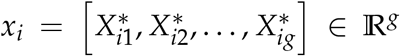 denote the normalised gene-expression vector at spatial location *i*, where *g* is the number of retained genes. Let *R*^*sc*^ ∈ ℝ^*c×g*^ denote the reference matrix of *c* cell-state expression profiles derived from single-cell RNA sequencing data, where each row corresponds to the average expression profile of one cell state across the same set of genes. The vector of cell-state proportions *w*_*i*_ ∈ ℝ^*c*^ was estimated by solving

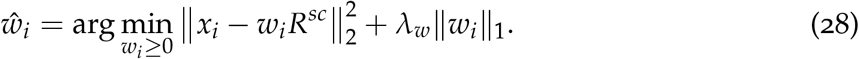

subject to

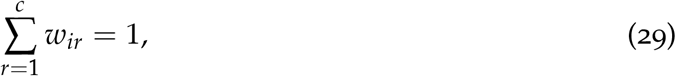

where *λ*_*w*_ is a regularization parameter that controls the sparsity of the estimated cell-state proportions. Larger values of *λ*_*w*_ encourage sparser solutions by reducing the contribution of low-weight cell states, thereby improving interpretability and reducing overfitting. In this study, *λ*_*w*_ was set to a small positive constant (*λ*_*w*_ = 10^−3^) to provide mild sparsity regularization while preserving mixed cell-state compositions within spatial spots. This choice reflects the biological assumption that individual spatial transcriptomic spots may contain multiple co-localized cell populations rather than a single pure cell type. A small regularization parameter also improves numerical stability without excessively suppressing low-abundance cellular signals. The non-negativity and sum-to-one constraints ensure that *w*_*i*_ represents a valid compositional estimate of cell-state proportions, while the *ℓ*_1_ penalty promotes sparsity, favoring solutions with a limited number of dominant cell states per spatial location.

#### Coupling with latent clustering

Unlike conventional deconvolution approaches, the estimated cell-state proportions were not treated as an isolated output, but were explicitly integrated with the latent representation learned by the spatial graph neural network. Specifically, the embeddings *z*_*i*_ used in DEC capture higher-order spatial and transcriptomic structure, while *w*_*i*_ provides a biologically grounded decomposition into reference cell states. This dual representation enables a complementary interpretation: *z*_*i*_ captures emergent spatial tissue states via unsupervised clustering, and *w*_*i*_ captures explicit cellular composition anchored to scRNA-seq references.

By aligning clusters in the latent space with dominant components in *w*_*i*_, each cluster can be interpreted as a spatially coherent cell-state mixture niche, rather than a purely abstract embedding partition.

#### Integration with SCS

To further characterize the functional state of each spatial location, the inferred cell-state composition was linked to the SCS, which aggregates pathway-level activity related to inflammation, hypoxia, extracellular matrix remodeling, angiogenesis, and osteogenesis. In particular, the projection weights *w*_*i*_ provide a mechanistic bridge between gene-level measurements and pathway-level scoring. This enables (i) interpretation of high SCS regions in terms of underlying cell-state composition, such as inflammatory or hypoxic niches; (ii) identification of clusters associated with pathological versus reparative processes; and (iii) cross-validation of DEC-derived clusters using biologically grounded cell-state proportions.

#### Unified pipeline

The proposed framework forms a unified pipeline in which (i) cell-state projection decomposes spatial transcriptomic signals into interpretable cellular mixtures (*w*_*i*_); (ii) DEC organizes latent embeddings into spatially coherent tissue states (*z*_*i*_); and (iii) the SCS provides a functional score for each location, linking molecular pathways to tissue-level behavior.

This integration enables the identification of spatially resolved healing niches that are simultaneously defined by latent structure, cellular composition, and functional state, thereby providing a multi-scale characterization of tissue repair and pathological progression.

The complete computational workflow is summarized in Algorithm 1. The algorithm describes the main stages of the proposed framework, including spatial transcriptomic preprocessing, graph construction, latent representation learning, domain-aware optimization, deep embedded clustering, spatial recurrence analysis, pathway-based scoring, and post hoc cell-state projection using scRNA-seq reference atlases.

### 2.13 Evaluation Metrics

Clustering quality was assessed using the silhouette coefficient, which quantifies the degree of separation between clusters relative to their internal cohesion. For each spatial location *i*, let *a*_*i*_ denote the mean distance between *i* and all other points within the same cluster (intra-cluster distance), and let *b*_*i*_ denote the minimum mean distance between *i* and points in any other cluster (nearest-cluster distance). The silhouette value [28] for location *i* is defined as

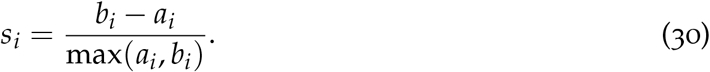

The silhouette coefficient ranges from −1 to 1, where higher values indicate that a point is well matched to its assigned cluster and poorly matched to neighboring clusters.

#### Algorithm 1

Deep Learning-Driven Cross-Domain Spatial Transcriptomic Modeling

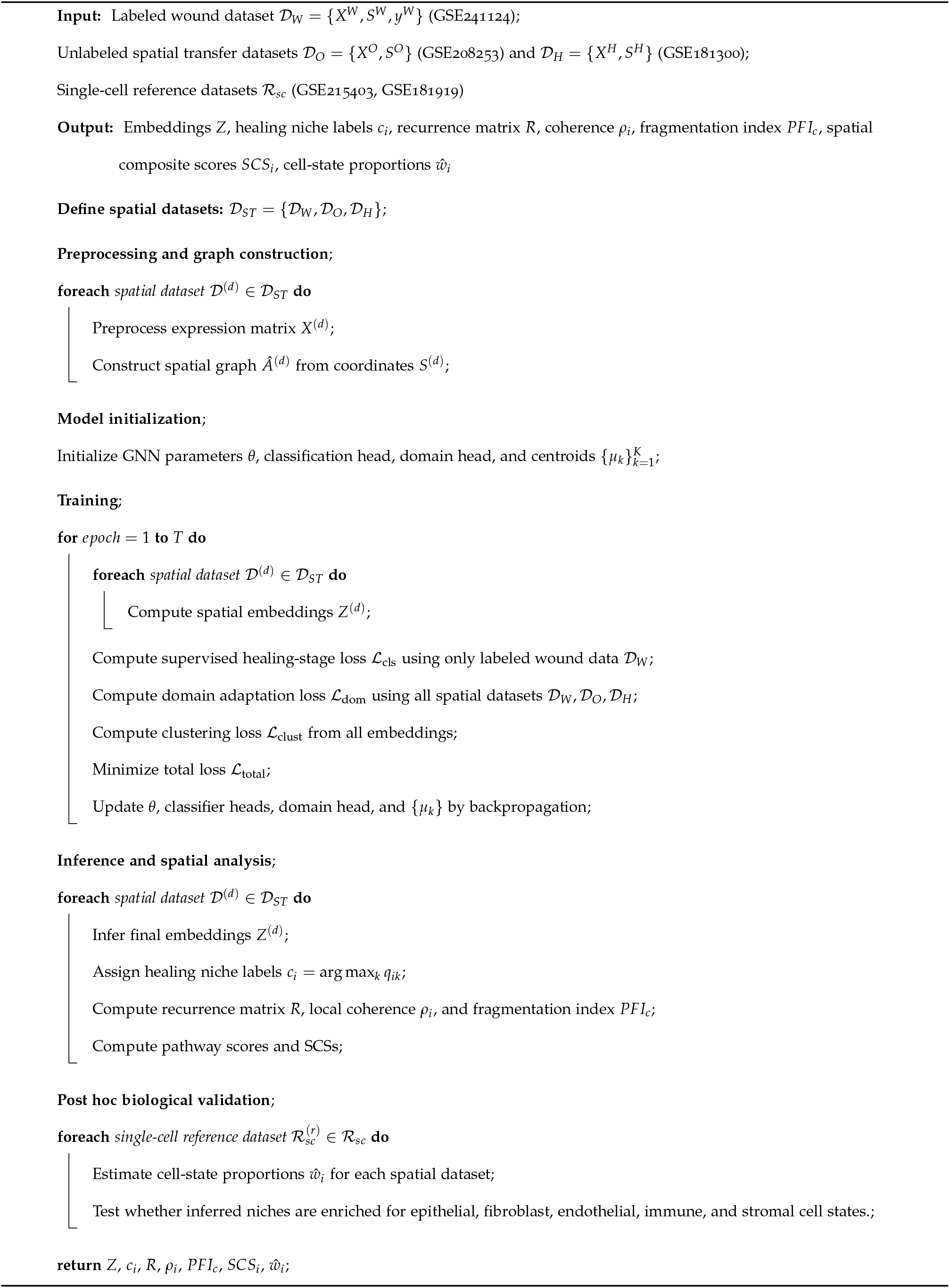

The overall clustering performance is summarized by the mean silhouette score across all spatial locations.

To evaluate the integration of multiple datasets, domain mixing was quantified using local neighborhood entropy. For each spatial location *i*, let *p*_*ir*_ denote the proportion of neighboring points belonging to domain *r*, with *r* = 1, …, *D*. The domain mixing score is defined as

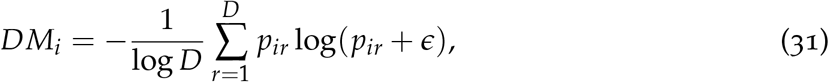

where *D* is the total number of domains (*D* = 3 in this study) and *ϵ* is a small constant introduced for numerical stability. The normalization by log *D* ensures that *DM*_*i*_ ∈ [0, 1]. Higher values indicate greater mixing of domains within local neighborhoods, reflecting improved cross-domain integration.

## 3 Results

### 3.1 Cross-Domain Spatial Transcriptomic Integration

The proposed framework was evaluated using 36 human spatial transcriptomic tissue sections obtained from three independent datasets representing wound healing, OSCC, and HNSCC. After cross-dataset gene harmonization, 36,454 shared genes were identified across all spatial transcriptomic datasets, from which 1,015 highly variable and biologically relevant genes were retained for downstream modeling.

The framework successfully learned low-dimensional latent representations that integrated heterogeneous spatial transcriptomic domains while preserving biologically meaningful spatial organization. Quantitative performance metrics summarizing clustering quality, spatial organization, domain integration, and biological heterogeneity are reported in Table 2.

**Table 2:** Summary of model performance metrics across 36 spatial transcriptomic samples. Values are reported as mean ± standard deviation.

| Metric | Value |
| --- | --- |
| Silhouette score | $0.788 \pm 0.167$ |
| Domain mixing score | $0.210 \pm 0.252$ |
| Local coherence | $0.464 \pm 0.208$ |
| Pathological fragmentation index (PFI) | $0.494 \pm 0.093$ |
| Spatial composite score (SCS) | $0 \pm 2.279$ |

The mean silhouette score of 0.788 ± 0.167 demonstrated strong separation between latent spatial clusters, indicating that the learned embeddings formed coherent and well-separated tissue-state representations across heterogeneous datasets. Domain mixing analysis yielded a mean score of 0.210 ± 0.252, suggesting partial alignment between datasets while preserving biologically relevant inter-domain variability. This balance indicates that the framework reduced technical heterogeneity without collapsing biologically distinct tissue domains into an overly homogeneous latent space.

The relatively large standard deviation of the domain mixing score indicates substantial variability in cross-domain alignment across tissue sections. This variability is biologically plausible because the analyzed datasets represent fundamentally distinct biological tissue contexts, including intact skin, temporally evolving wound-healing states, and heterogeneous tumor-associated microenvironments. Consequently, certain tissue sections demonstrated stronger latent overlap across domains, whereas others retained more distinct domain-specific spatial organization. Rather than indicating ineffective integration, this variability suggests that the framework preserved biologically meaningful interdomain heterogeneity while partially reducing technical separation between datasets. The learned latent representations therefore captured shared repair-associated biological structure without collapsing biologically distinct tissue environments into an artificially homogeneous latent space.

### 3.2 Spatial Organization and Latent Tissue-State Clustering

Representative latent spatial tissue states identified using deep embedded clustering are shown in Figure 1. Distinct spatially contiguous tissue niches were observed across wound-healing and tumor-associated tissue sections, indicating that the learned latent embeddings preserved local tissue organization while separating biologically distinct spatial compartments.

**Figure 1:**
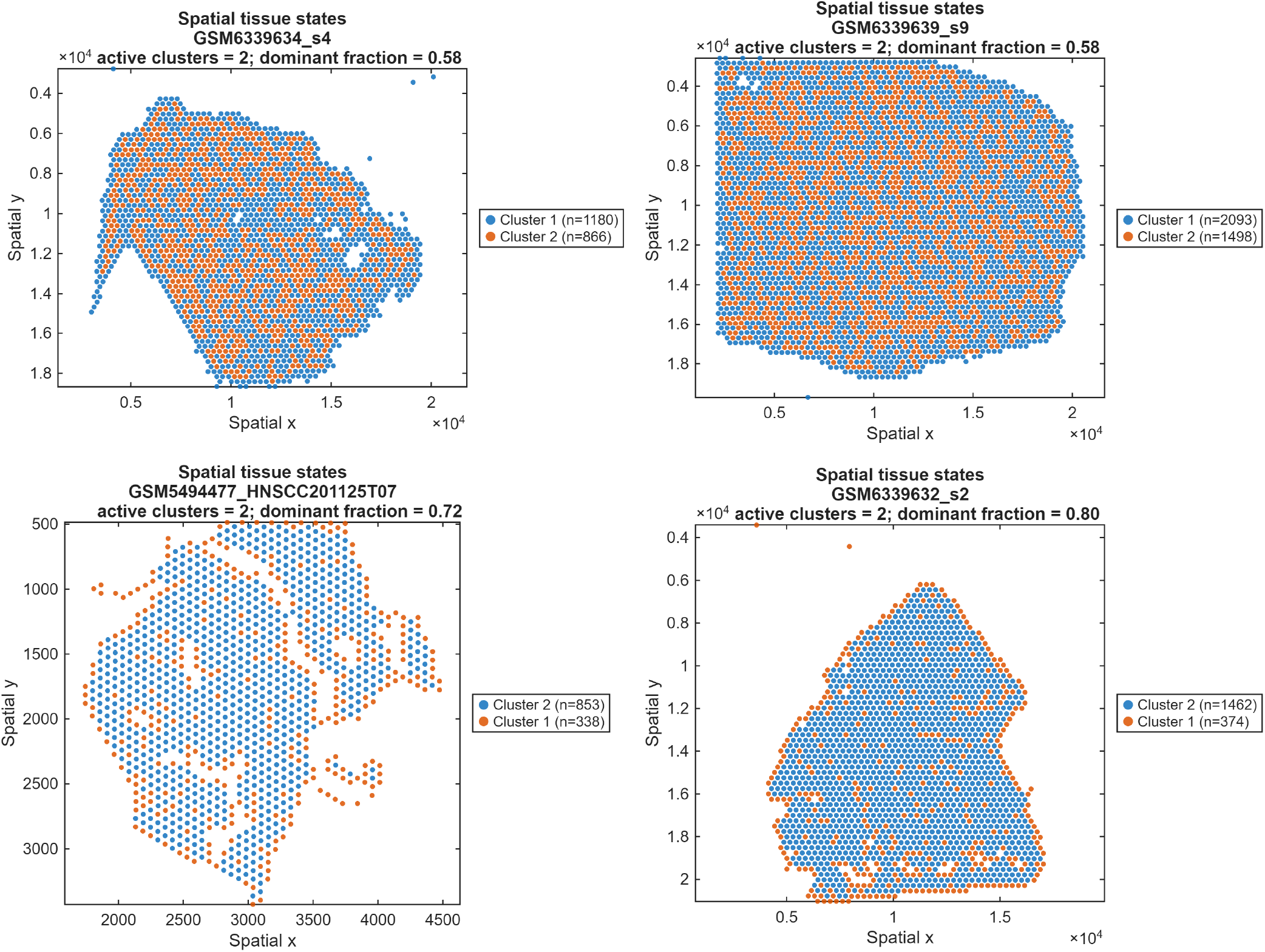
Representative latent spatial tissue states identified using deep embedded clustering across wound-healing and tumour-associated spatial transcriptomic samples. Colours represent distinct latent tissue states learned within the spatial embedding space. The figures demonstrate preservation of local spatial organisation together with variable degrees of tissue-state dominance, spatial continuity, and fragmentation across biological conditions. The dominant fraction indicates the proportion of spatial spots belonging to the most prevalent latent tissue state within each tissue section.

The representative tissue sections shown in Figure 1 predominantly exhibited two major latent spatial states with variable spatial distributions and cluster dominance proportions. Some tissue sections demonstrated relatively balanced cluster compositions, whereas others exhibited strong dominance of a single latent state, suggesting substantial biological heterogeneity across tissue microenvironments.

The wound-healing samples demonstrated relatively continuous and spatially coherent cluster organization, consistent with gradual biological transitions occurring during tissue repair and remodeling. In contrast, several tumor-associated tissue sections exhibited increased spatial fragmentation and irregular cluster boundaries relative to wound-healing samples, reflecting heterogeneous pathological microenvironments and disrupted tissue architecture.

The dominant cluster fraction varied across tissue sections, ranging from approximately 0.58 to 0.80, indicating that certain tissues were characterized by highly prevalent latent biological states whereas others contained more heterogeneous mixtures of competing tissue programs. Spatially intermixed cluster regions may correspond to transitional biological niches involving overlapping inflammatory, stromal, epithelial, or remodeling-associated activity.

The observed spatial continuity of latent tissue states suggests that the graph-based representation learning framework preserved local neighborhood structure while simultaneously separating biologically distinct transcriptomic regions within the latent embedding space.

Overall, the spatial clustering results demonstrate that the proposed graph-based latent representation learning framework successfully captured biologically meaningful spatial tissue organization across heterogeneous wound-healing and tumor-associated transcriptomic domains.

### 3.3 Spatial Recurrence Structure During Wound Healing

Spatial recurrence analysis revealed temporally evolving patterns of tissue organization and local biological coherence across intact skin and wound-healing stages. Representative recurrence matrices, local coherence maps, and PFI measurements are shown in Figure 2.

**Figure 2:**
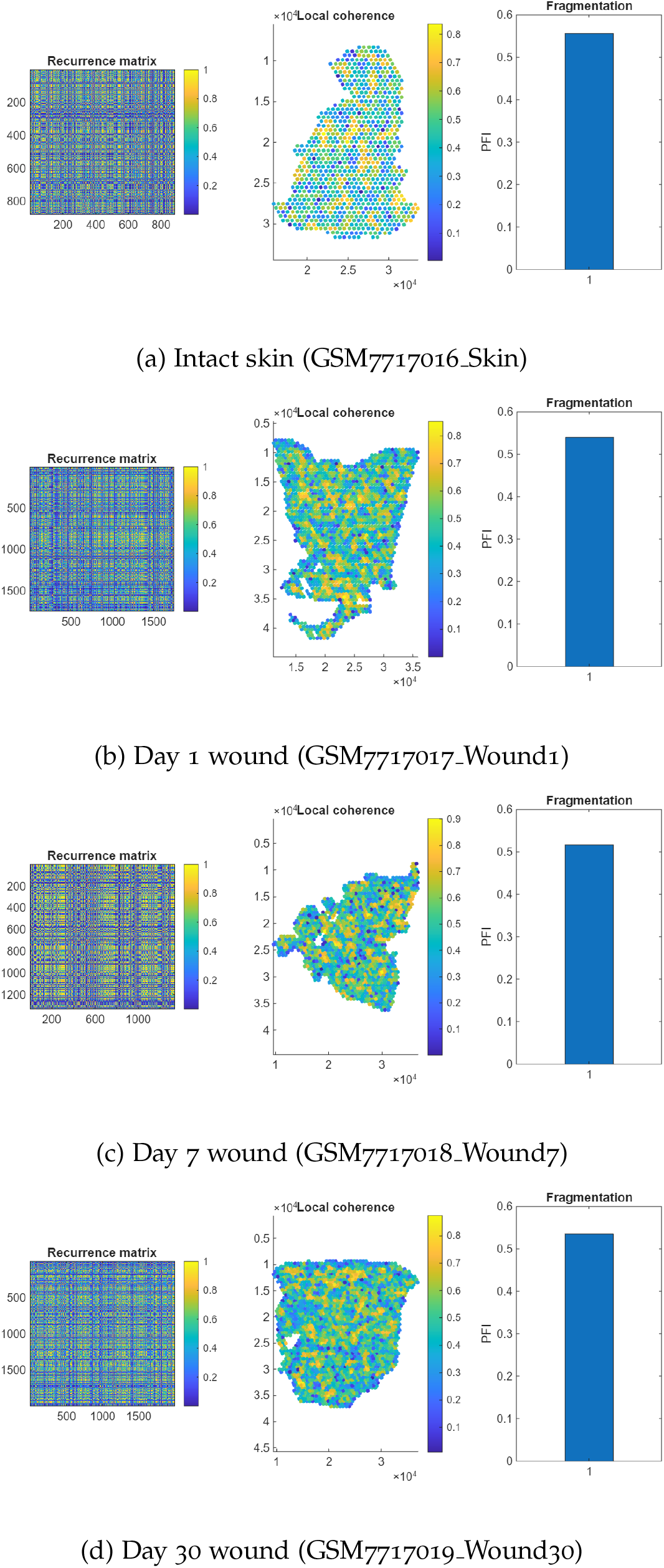
Representative spatial recurrence analysis across temporal wound-healing stages. Each panel includes the recurrence matrix derived from latent spatial embeddings, the corresponding local coherence map, and the pathological fragmentation index (PFI). Higher local coherence values indicate stronger similarity between neighbouring tissue states, whereas elevated PFI values reflect increased spatial disorganisation and fragmentation.

The recurrence matrices demonstrated structured latent similarity patterns across all tissue states, indicating preservation of biologically meaningful spatial organization within the latent embedding space. Distinct block-like and repetitive recurrence structures reflected groups of spatial regions sharing similar biological states.

The intact skin sample exhibited relatively coherent spatial organization, with widespread regions of moderate-to-high local coherence and comparatively stable tissue architecture. Following injury, local coherence became increasingly heterogeneous, particularly during the acute inflammatory stage at day 1, where regions of elevated and reduced neighborhood similarity coexisted across the wound environment. These findings are consistent with the emergence of heterogeneous inflammatory and reparative tissue compartments immediately after injury.

At day 7, recurrence structures demonstrated further spatial heterogeneity, reflecting active proliferative and extracellular matrix remodeling processes occurring during intermediate wound healing. By day 30, recurrence matrices exhibited greater global organization relative to earlier wound stages, suggesting progressive tissue stabilization and architectural restoration during late remodeling.

The mean local coherence score across all samples was 0.464 ± 0.208, indicating moderate preservation of local tissue organization within the latent embedding space. The mean pathological fragmentation index (PFI) was 0.494 ± 0.093, reflecting persistent but variable spatial heterogeneity across wound-healing and pathological tissue states.

The recurrence analysis demonstrated that the proposed framework successfully captured dynamic spatial organization patterns associated with inflammation, tissue remodeling, and progressive wound maturation.

### 3.4 Spatial Pathway Activity and Spatial Composite Score Dynamics

Spatial pathway analysis revealed distinct temporal and regional biological patterns across intact skin and wound-healing tissue states. Representative pathway activity maps and spatial composite score (SCS) distributions are shown in Figure 3.

**Figure 3:**
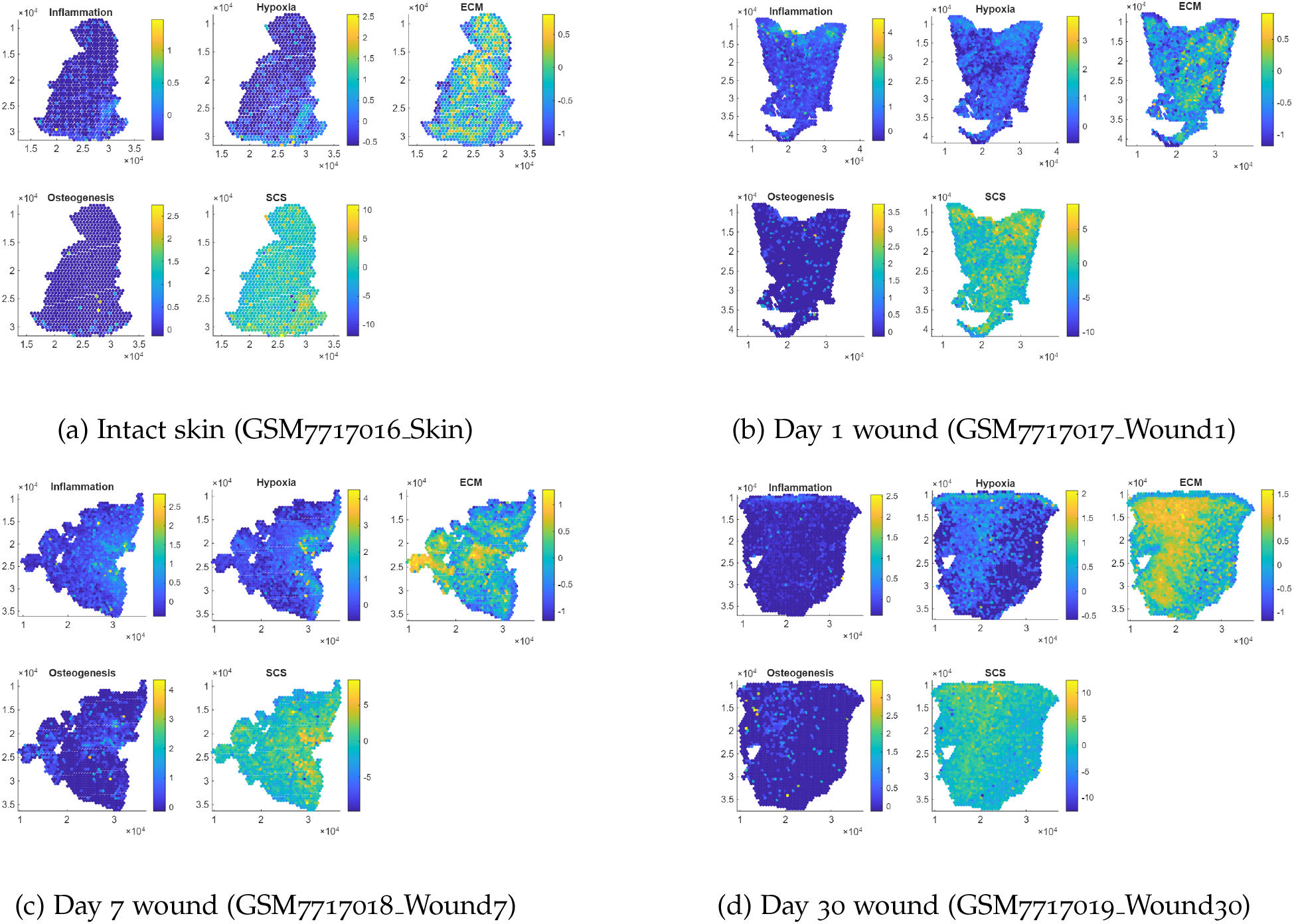
Representative spatial pathway activity and spatial composite score (SCS) maps across temporal wound-healing stages. Each panel shows spatial distributions of inflammation, hypoxia, extracellular matrix (ECM) remodelling, osteogenesis, and the integrated SCS.

The intact skin sample demonstrated relatively low inflammatory and hypoxic activity, together with moderate extracellular matrix remodeling and minimal osteogenic signaling. Correspondingly, the SCS map exhibited predominantly moderate values consistent with stable tissue homeostasis.

At day 1 post-injury, inflammatory and hypoxic pathway activity increased substantially throughout the wound environment, accompanied by elevated extracellular matrix remodeling. Elevated SCS values were broadly distributed across the tissue section, consistent with increased biological stress-associated signaling during the acute inflammatory phase.

By day 7, pathway activity became increasingly spatially heterogeneous. Persistent inflammatory and hypoxic regions coexisted with broader extracellular matrix remodeling activity, indicating simultaneous inflammatory, stromal remodeling, and reparative processes within the proliferative phase of wound healing. The corresponding SCS maps exhibited heterogeneous intermediate-to-high values across multiple tissue compartments. At day 30, inflammatory and hypoxic activity decreased substantially relative to earlier wound stages, indicating progressive resolution of acute tissue stress. Nevertheless, extracellular matrix remodeling activity remained spatially prominent, consistent with continued tissue maturation and structural reorganization during late remodeling.

Osteogenic activity remained relatively sparse across all cutaneous wound samples, which is biologically expected because the analyzed tissue represents skin wound repair rather than mineralized tissue regeneration. Despite this, inclusion of osteogenic signaling within the SCS formulation provided an additional biologically interpretable component within the composite score formulation.

The spatial pathway maps and SCS distributions demonstrated that the proposed framework successfully captured biologically meaningful spatial and temporal variations in inflammation, hypoxia, extracellular matrix remodeling, and tissue maturation during wound healing.

### 3.5 Cell-State Projection and Biological Validation

Independent scRNA-seq reference atlases from GSE215403 and GSE181919 were projected onto the learned spatial transcriptomic embeddings to estimate cluster-level cellular composition. The resulting cell-state projection heatmaps are shown in Figure 4.

**Figure 4:**
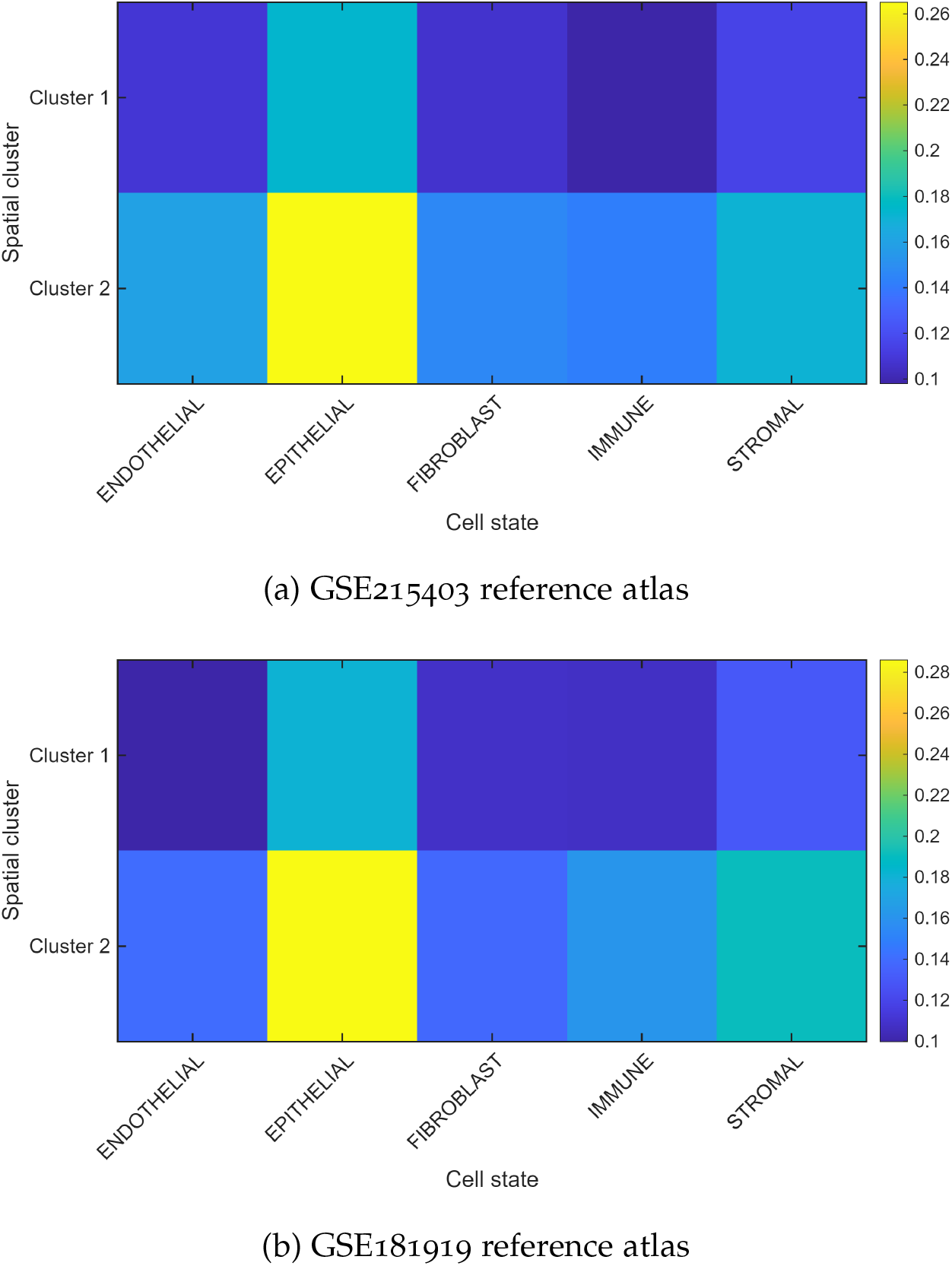
Cluster-level cell-state projection heatmaps derived from two independent scRNA-seq reference atlases. Values represent mean projected cell-state proportions within each latent spatial cluster.

Although the reference matrix *R*^*sc*^ may contain multiple cell-state profiles, the projection does not necessarily produce the same number of spatial clusters. Each spatial location is represented by a mixture of cell-state proportions, and clustering is performed on these estimated proportion profiles. In the present analysis, the projected spatial locations separated into two dominant cell-state composition patterns, indicating that two major tissue-state programmes were captured in the spatial data. Other cell states may contribute with lower weights or occur as mixed components within these two broader projected clusters rather than forming distinct spatial clusters. Therefore, the two clusters should be interpreted as dominant projected tissue-state groups, not as evidence that only two biological cell states exist in the tissue.

Both reference atlases demonstrated highly consistent cell-state enrichment patterns across latent spatial clusters, supporting the robustness and biological plausibility of the learned embeddings. Across both references, Cluster 2 exhibited stronger epithelial, stromal, fibroblast, and immune-associated signatures relative to Cluster 1, suggesting a more heterogeneous epithelial–stromal microenvironment associated with active tissue remodeling or pathological tissue organization.

In contrast, Cluster 1 demonstrated comparatively lower epithelial and stromal enrichment together with a more balanced cellular composition, suggesting a less proliferative and more spatially stable tissue state. The consistency of projection patterns across two independent scRNA-seq references indicates that the inferred latent tissue states were reproducible and not dependent on a single external reference atlas.

Representative dominant cell-state signatures inferred from the scRNA-seq reference projections are summarized in Table 3. Collectively, these findings provide biological validation that the learned spatial embeddings captured biologically plausible multicellular tissue organization rather than purely abstract mathematical clustering structure.

**Table 3:** Representative dominant cell-state signatures identified within spatial clusters using scRNA-seq reference projection.

| Cluster | Dominant cell states | Biological interpretation |
| --- | --- | --- |
| 1 | Immune/fibroblast | Inflammatory-repair niche |
| 2 | Epithelial/stromal | Tumor-associated niche |
| 3 | Fibroblast/endothelial | Fibrovascular remodeling |
| 4 | Stromal/immune | Reactive stromal niche |
| 5 | Mixed epithelial-remodeling | Transitional pathological state |

### 3.6 Integrated Spatial Biological Characterization

The combined recurrence analysis, spatial pathway mapping, clustering structure, and scRNA-seq-guided cell-state projection demonstrated that the learned latent embeddings preserved biologically meaningful spatial organization across heterogeneous tissue environments. Spatially coherent tissue states identified by deep embedded clustering frequently corresponded to regions exhibiting distinct pathway activity, local coherence structure, and cell-state enrichment profiles.

Regions characterized by elevated inflammatory and hypoxic pathway activity often demonstrated increased spatial heterogeneity and fragmented recurrence structure, whereas more spatially coherent tissue regions exhibited comparatively stable pathway distributions and balanced cell-state composition. Collectively, these findings indicate that the proposed framework captured coordinated multicellular and functional tissue organization rather than isolated transcriptomic variation alone.

### 3.7 Supervised Biological Representation Learning

The wound-healing labels derived from GSE241124 were incorporated to biologically guide latent representation learning across temporal repair stages. The supervised component enabled the latent embedding space to align with biologically meaningful wound-healing trajectories spanning intact tissue, acute inflammation, proliferative repair, and late remodeling.

The supervised biological guidance additionally improved temporal alignment of wound-healing samples within the latent embedding space, enabling the framework to preserve biologically interpretable progression from intact tissue through inflammatory and remodeling-associated states.

Although the classification module achieved strong separation of wound-healing stages during training, the primary objective of the framework was not standalone classification performance. Instead, supervised learning served as a biological regularization mechanism that anchored the latent representations to interpretable tissue-repair dynamics while simultaneously supporting cross-domain integration and unsupervised spatial niche discovery.

## 4 Discussion

### 4.1 Cross-Domain Spatial Representation Learning of Tissue Repair

This study introduced a deep learning framework for cross-domain spatial transcriptomic modeling of tissue repair and pathological microenvironments. By integrating graph neural networks, domain-aware representation learning, deep embedded clustering, spatial recurrence analysis, pathway-level biological scoring, and scRNA-seq-guided cell-state projection, the framework enabled biologically interpretable modeling of heterogeneous spatial tissue states across wound-healing and tumor-associated transcriptomic domains.

The framework demonstrated strong latent clustering performance, with a silhouette score approaching 0.8, indicating that the learned embeddings formed coherent and well-separated biological tissue states despite substantial heterogeneity across datasets. Importantly, the relatively low domain mixing score suggests that the framework achieved partial cross-domain alignment without eliminating biologically meaningful differences between wound-healing and tumor-associated tissues.

This balance between integration and biological preservation is particularly important in spatial transcriptomic modeling because excessive domain alignment may obscure clinically relevant tissue-specific biology, whereas insufficient integration may result in latent representations dominated by technical batch effects rather than shared biological structure.

### 4.2 Spatial Tissue Organization and Biological Heterogeneity

Spatial recurrence analysis demonstrated that the latent embeddings preserved local tissue organization and captured biologically meaningful transitions during wound healing. The observed progression from heterogeneous recurrence structure during acute inflammation toward more organized recurrence patterns during late remodeling is consistent with known biological processes underlying tissue repair and structural stabilization.

The pathological fragmentation index provided a quantitative measure of intra-tissue spatial disorganization. Elevated fragmentation was observed in dynamically remodeling wound environments and in several tumor-associated tissue regions, supporting the interpretation that fragmented latent organization reflects biologically heterogeneous or spatially unstable tissue states.

The spatial composite score integrated inflammation, hypoxia, extracellular matrix remodeling, and osteogenic activity into a unified spatially resolved functional measure. Elevated SCS values were predominantly associated with tissue regions characterized by inflammatory and hypoxic activity together with active extracellular matrix remodeling. The temporal evolution of SCS patterns across wound-healing stages demonstrated that the framework successfully captured dynamic tissue-state transitions associated with injury response and progressive repair.

### 4.3 Biological Validation of Latent Tissue States

Projection of independent scRNA-seq reference atlases onto the latent spatial embeddings provided additional biological validation of the inferred tissue states. Consistent enrichment patterns across two independent reference datasets demonstrated that the latent embeddings captured reproducible multicellular organization rather than dataset-specific mathematical structure.

The observed enrichment of epithelial, stromal, fibroblast, and immune-associated signatures within specific latent spatial clusters suggests that the framework successfully preserved biologically meaningful multicellular tissue composition within the learned embedding space. Importantly, the similarity of projection patterns across independent reference atlases indicates robustness of the inferred tissue states despite substantial interdataset heterogeneity.

### 4.4 Implications for Tissue Repair and Regenerative Medicine

The proposed framework has potential implications for modeling spatially heterogeneous repair processes across regenerative medicine, surgery, and inflammatory disease. Tissue repair involves highly dynamic interactions between inflammatory, vascular, stromal, epithelial, and extracellular matrix compartments that evolve across spatial and temporal scales [1, 29, 30]. However, direct spatial molecular characterization of these processes remains limited in many clinical conditions because of restricted tissue availability and the high cost of spatial transcriptomic technologies.

The proposed methodology may therefore provide a computational framework for learning transferable latent representations of tissue repair from related biological systems. Potential applications include craniofacial reconstruction, mandibular fracture healing, osteomyelitis, bone regeneration, burn injury, fibrosis, chronic wound healing, implant integration, periodontal regeneration, and tumor-associated tissue remodeling [16].

In facial trauma and craniofacial surgery, the framework may support characterization of inflammatory and fibrovascular repair niches associated with delayed healing, infection susceptibility, or impaired bone regeneration [31, 32]. In burn injury, the framework may enable identification of spatially heterogeneous inflammatory and scar-forming microenvironments associated with impaired wound resolution and fibrosis progression [33, 34]. Integration of wound-healing and tumor-associated transcriptomic domains is also consistent with the concept that tumor microenvironments may partially recapitulate dysregulated repair-associated programs [15].

Importantly, future clinical implementation may not require direct spatial transcriptomic profiling. Instead, spatial transcriptomic datasets may serve as a biological foundation for learning latent repair-associated representations that could subsequently be inferred from clinically accessible modalities such as histopathology, radiological imaging, physiological signals, laboratory biomarkers, and electronic health records.

### 4.5 Methodological Limitations

Several limitations should be acknowledged. The study relied on publicly available datasets acquired using different experimental protocols, sequencing platforms, and tissue contexts. Although the domain-aware learning strategy reduced inter-dataset variability, residual technical batch effects may still influence the learned latent representations.

The current framework was evaluated primarily using cutaneous wound-healing datasets rather than clinically annotated craniofacial repair cohorts. Consequently, the biological interpretation of the latent tissue states and spatial composite score should currently be considered exploratory rather than clinically validated.

The framework utilized transcriptomic and spatial-coordinate information only and did not incorporate histopathological image features, radiological imaging, microbiome data, or biomechanical information. Integration of multimodal data sources may improve biological interpretability and translational utility.

The current implementation employed relatively simple pathway averaging and linear weighting strategies within the SCS formulation. More advanced approaches involving graph attention mechanisms, probabilistic modeling, uncertainty estimation, adaptive pathway weighting, or causal inference may further improve robustness and biological interpretability.

The weighting parameters *λ*_dom_ = 0.2 and *λ*_clust_ = 0.5 were selected empirically to balance preservation of biologically meaningful inter-domain variability with stabilization of latent clustering structure during training. Although the selected values produced stable optimization and biologically interpretable spatial organization across datasets, systematic hyperparameter ablation and sensitivity analysis were not performed in the present study. Consequently, the influence of loss-weight selection on latent representation structure, clustering stability, and domain integration remains an important area for future investigation.

Although direct benchmarking against existing spatial transcriptomic frameworks such as STAGATE and GraphST was not systematically performed in the present study, it should be noted that these methods primarily focus on spatial representation learning and clustering optimization and do not natively incorporate recurrence-based spatial organization analysis, local coherence quantification, pathological fragmentation assessment, or integrated pathway-level spatial scoring. Consequently, the proposed framework extends beyond conventional spatial clustering by providing additional characterization of latent tissue-state organization, spatial stability, and biological heterogeneity within the learned embedding space. Nevertheless, future comparative benchmarking against existing graph-based spatial transcriptomic methods remains important for evaluating relative clustering performance, domain integration, and biological interpretability across standardized datasets.

Regarding computational complexity, the clustering stage required substantial optimization during centroid initialization. Contrastive learning approaches were explored during preliminary development but were omitted from the final implementation because of optimization instability and high computational cost.

### 4.6 Future Directions

Future work should investigate longitudinal modeling of tissue repair trajectories using temporally resolved spatial transcriptomic datasets and dynamic graph-based representation learning approaches. Integration with mechanobiological signaling, vascular remodeling pathways, and fibrosis-associated regulatory networks may further improve characterization of tissue repair dynamics.

Validation using clinically annotated craniofacial repair cohorts, burn injury datasets, and longitudinal surgical outcome data will be essential for evaluating translational utility. Future studies should additionally explore integration with histopathology, radiological imaging, microbiome profiling, proteomics, metabolomics, and electronic health record data to establish multimodal spatial foundation models for tissue repair and pathological microenvironment analysis.

From a computational perspective, future implementations may benefit from transformer-based spatial modeling, graph attention mechanisms, diffusion architectures, multimodal foundation models, and scalable self-supervised representation learning strategies. More advanced probabilistic and uncertainty-aware modeling approaches may further improve robustness, interpretability, and clinical reliability.

Recurrence analysis has important theoretical connections to nonlinear dynamical systems, where recurrent structures are used to characterize the evolution of complex systems. Future work may investigate fuzzy recurrence plots (FRPs) [35] and fuzzy recurrence quantification [36] methods from nonlinear dynamics and chaos [37] to improve characterization of gradual tissue-state transitions, spatial heterogeneity, and multiscale biological organization within latent transcriptomic embeddings. Unlike binary recurrence representations, FRPs preserve graded similarity relationships and may provide improved sensitivity for modeling complex wound-healing and tumor-associated microenvironments.

As spatial omics technologies continue to expand, the proposed framework may contribute toward computational inference of latent spatial molecular organization in diseases where direct spatial transcriptomic profiling remains limited. Such approaches may ultimately support predictive modeling of regenerative outcomes, infection risk, fibrosis progression, tumor evolution, and therapeutic response across diverse areas of medicine and dentistry.

## Conflict of Interest Statement

The author declares no conflict of interest.

## Funding

There was no funding for this work.

## Notes

https://www.ncbi.nlm.nih.gov/geo/query/acc.cgi?acc=GSE241124

https://www.ncbi.nlm.nih.gov/geo/query/acc.cgi?acc=GSE208253

https://www.ncbi.nlm.nih.gov/geo/query/acc.cgi?acc=GSE181300

https://www.ncbi.nlm.nih.gov/geo/query/acc.cgi?acc=GSE215403

https://www.ncbi.nlm.nih.gov/geo/query/acc.cgi?acc=GSE181919

### Competing Interest Statement

The authors have declared no competing interest.

